# Transfer of Learned Object Manipulations between Two- and Five-Digit Grasps

**DOI:** 10.1101/2025.02.24.639552

**Authors:** Jordana Ulloa-Marquez, Jennifer Gutterman, Marco Santello, Andrew M. Gordon

## Abstract

Successful object manipulation involves integrating object properties into a motor plan and scaling fingertip forces through learning. This study investigated whether learned manipulations using a two-digit grip transfer to a five-digit grip and vice versa, focusing on the challenges posed by added degrees of freedom in force distribution. The goal of the task was to exert the necessary compensatory torque (Tcom) and vertical forces to minimize object roll on a visually symmetrical object that with an asymmetrical mass distribution. To examine this, subjects performed blocked consecutive learning trials before switching grip type. Our results support the learning transfer between two-digits and five-digit grasp configurations despite challenges in maintaining perfect stability during the grip switch. Subjects adapted their grip forces (GF), center of pressure (CoP), and Tcom to minimize object roll, with significant improvements observed from novel (1^st^) to transfer (11^th^) trials. These findings suggest high-level, effector-independent representations of object manipulation that enable generalization across grip types, though some limitations in force distribution and digit position arise during transfers.

## INTRODUCTION

In order to successfully grasp or manipulate an object, its physical properties (e.g., size, texture, and shape of object) are integrated into a motor plan well before the hand touches the object (e.g., Jeannerod et al., 1995; van Polanen and Davare, 2015). Repeated lifting experience helps refine this complex sequence of movements through the acquired implicit knowledge needed to learn how to lift objects with the appropriate level of forces for the object’s weight and scale the fingertip forces accordingly (Fu and Santello, 2014; Gordon et al., 1991; Westling and Johansson, 1984).

When we lift an object with an asymmetrical weight distribution, it requires the digits to exert a net vertical force as well as a torque to prevent the object from rolling (which is a rotation about the x-axis) (Fu et al., 2011). This is accomplished by developing a compensatory torque (Tcom) to counteract the external object torque determined by the object’s center of mass (CM) location prior to lift-off (Fu et al., 2010, 2011; Fu and Santello, 2014; Lee-Miller et al., 2021). When subjects’ grasp is not constrained by the location of the force sensors (i.e., subjects are free to choose their grasp location on the object’s vertical surface), they also place the digit on the CM side of the objects higher than the opposing digit, to create a passive torque (J. Lukos et al., 2007; J. R. Lukos et al., 2008). Over repeated lifts subjects learn to modulate digit forces as a function of digit placement to exert a compensatory torque in an anticipatory manner, and this coordination between digit forces and positions is critical for successful manipulation (Fu et al., 2010, Fu et al., 2011; Lee-Miller et al., 2021). Depending on the necessary Tcom required for the lift, the digit placement and digit forces may vary from trial to trial.

Previous precision grasp studies (e.g., (Fu et al., 2011; Lee-Miller et al., 2021) have investigated the generalizability of these motor representations, i.e., Tcom, and raise the question of the extent of motor equivalence in grasping: Are these representations of grasping actions specific to and/or constrained by the effectors that we use to learn the given action? Or are they independent from how they are learned? When subjects grasp objects at constrained contacts and manipulate them with a given object texture or mass, they are able to transfer the information about these object properties for anticipatory control of digit forces (i.e., before feedback becomes available) during subsequent lifts with the contralateral hand (Gordon et al., 1994; Gordon and Salimi, 2004; Westling and Johansson, 1987). For objects with an asymmetric CM, if the object or subject is moved or rotated in between lifts, subjects maintain appropriate anticipatory force scaling as long as the relationship between the subject’s specific digit (thumb or index finger) and the CM distribution remains the same (Salimi et al., 2000). For such objects, subjects are seamlessly able to transfer information related to the CM location during unconstrained lifts with a two-digit precision grip and subsequently transfer this information during lifts with three-digits (i.e., adding the middle finger) and vice-versa (Fu et al., 2011). Subjects were able to match the Tcom used for the pre-transfer grip configuration following transfer to the other grip configuration, and thus there was not a greater roll on these trials. Interestingly, subjects used different digit force-position coordination patterns in the two grip types (Fu et al., 2011). Together, these findings suggest a high-level (task) representation of the learned object manipulation (i.e., Tcom), rather than effector-level representation (i.e., the specific digit positions or forces that must be used). The addition or removal of the adjacent middle finger is likely a relatively minor perturbation since the close proximity of these fingers may easily involve utilization of a “virtual finger” (e.g., Arbib et al., 1985; Sun et al., 2011) with the center of pressure simply shared between the two digits. Larger perturbations such as rotating objects with an asymmetric CM between lifts (Zhang et al., 2010) initially result in failed transfer (i.e., large rolls). Subjects initially fail to generate a compensatory moment to counter the external moment caused by the new CM location. However, after several object rotations subjects reduced object roll on the initial post-rotation trials by anticipating the new CM location through the modulation of digit placement but not fingertip forces. This further suggests that transfer of digit placement and forces may differ.

The aim of the current study was to further investigate whether these learned grasping manipulations are transferrable, or if they are constrained to the learning conditions of these manipulations. Our primary aim was to examine the possible constraints of these grasping internal representations on learning transfer after adding or removing digits. We tested transfer of learned manipulations using a two-digit precision grip to whole hand grasping, and vice-versa. Given the large number of degrees of freedom in the force sharing patterns across all four digits opposing the thumb during whole hand grasping, the complexity of choosing among the many solutions in force distribution to create a Tcom increases immensely. Thus, the previous interpretation of successful transfer with a smaller number of degrees of freedom (i.e., from 2 to 3, and 3 to 2 digits; Fu et al. 2011) may not be a general property of the CNS. Specifically, how many degrees of freedom are added or removed might impact whether learned manipulation can or cannot be transferred. While adding more degrees of freedom introduces many possible solutions for digit force distribution, we hypothesize that subjects will use a high-level (task) representation of the learned object manipulation, such as maintaining the object’s stability and controlling its orientation, rather than effector-specific force sharing patterns and CoP transfer. We predict that subjects would effectively demonstrate within-hand transfer of learned manipulation between two-digit and five-digit grasping. We expect the Tcom in the pre-transfer trial to be significantly higher than in the novel trial (i.e., 1^st^ trial where subjects cannot visually assess object CM), with little difference between the pre-transfer and transfer trial Tcoms, nor between the transfer and final trials. Successful transfer will be indicated by a significant improvement in Tcom (i.e., the anticipatory torque approaching the magnitude required to match external torque) from the novel trial to the transfer trial, accompanied by minimized roll (i.e., smaller roll angles approaching 0 degrees).

## MATERIALS AND METHODS

### Subjects

20 right-hand dominant (9 males and 11 females; mean age and SD: 25.21 +/-4.13 years) participated in this study. Subjects reported no previous history of orthopedic, neurological trauma, or pathology of the upper limbs and were naïve to the purpose of the study. Informed written consent was obtained prior to participation in compliance with the Declaration of Helsinki. The study was approved by the Teachers College, Columbia University Institutional Review Board.

### Apparatus

A custom-made grip device (adapted from Fu et al., 2011) was used to measure the digit forces and the center of pressure (See Fig. 1A). The device was composed of two 6-axis force/torque transducers (Mini 40, ATI Industrial Automation, NC, USA) that were mounted collinearly onto the grip device on both the thumb side and fingers separately. The force transducers measured both grip and load forces, as well as the torque exerted on the device (with a resolution of 0.02 N, 0.01 N, and 0.125 Nmm respectively). An electromagnetic sensor (Polhemus Fastrak, 0.005 mm range, 0.025° resolution) was attached on top of the device to measure the vertical distance and object roll. The grip surfaces were comprised of two carbon fiber surfaces (dimensions: 14 cm x 4.5 cm) and mounted vertically onto each F/T transducer. The distance (grip width) between the surfaces was 6 cm. The grip device was designed as an inverted T-stand made of plexiglass to allow for the change in mass distribution to the left, middle, or right of the device midline by inserting a mass (400 g) into one or both compartments. Cardboard covers concealed the mass compartments on the bottom of the device to prevent any visual identification of the object’s center of mass (Fig. 1A). The overall weight of the device was 880 g and the calculated target object Tcom was 20.67 Ncm.

**Fig 1.**
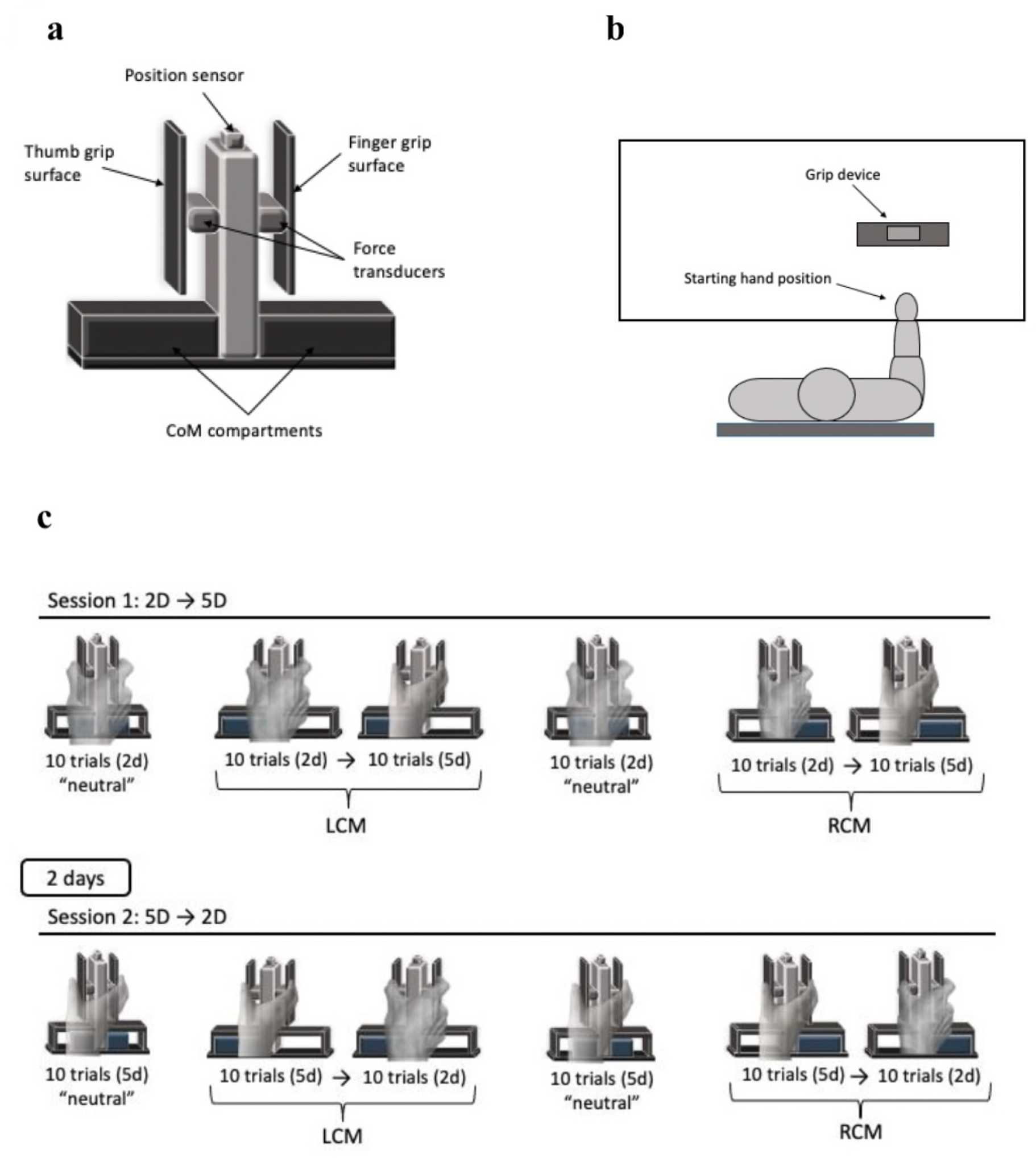
Experimental Apparatus. **(a)** Physical appearance of the grasp object including the covers to conceal the CM compartments to the subjects. **(b)** Aerial view of the setup. **(c)** Example of one of the two trial conditions associated with switching between two to five digits (2d → 5d) and vice versa (5d → 2d). Subjects were tested on experimental session 2, 2 days after experimental session 1. Trial sequences were counterbalanced across subjects. All Subjects began each experimental session completing the L_CM_ using both grip types, followed by the same number of trials for the R_CM._

### Experimental procedures

Subjects were asked to sit facing the grip device with their right elbow flexed (approximately 90° in the parasagittal plane), align their shoulder to the midline of the device, and place their pronated right hand on the edge of the table (Fig. 1B). After the auditory cue was given, subjects reached and grasped the object anywhere on the grip surfaces with the tip of their thumb and finger(s) for the respective side and lifted the grip device at a natural self-selected speed to be adjacent to a 10 cm marker. They were instructed to hold it in the air until they heard a second auditory cue 1 s after the object reached the specified height, at which point they replaced it back to its start location. Subjects were asked to only use their thumb and index finger to contact the grip surface for the 2-digit (*2D*) grasp condition and all digits for the 5-digit (*5D*) condition. Compliance was visually verified by one of the experimenters. The experimenter instructed the participant to maintain a self-selected natural pace and emphasized the goal of the task to keep the object as steady as possible.

Each participant performed the task under the following experimental conditions which were conducted in two sessions with a two-day break in between: two-digit grasp (thumb and index finger; *2D*) and five-digit grasp (thumb, index, middle, third, and little finger; *5D*). The two conditions (*2 Digit* → *5 Digit* and *5 Digit* → *2 Digit*) were counterbalanced among the subjects. Ten subjects completed the transfer sequence from *2 Digit* → *5 Digit* and the other ten completed the transfer from *5 Digit* → *2 Digit*. Each session was divided into four blocks to include a practice block, both grasp conditions and the different center of mass (CM) locations (*2 Digit* → *5 Digit* (L_CM_), *2 Digit→ 5 D* (R_CM_), *5 Digit* → *2 Digit* (L_CM_), and *5 Digit* → *2 Digit* (R_CM_)), with a between-condition “neutral” block to “wash out” the prior lifts with the asymmetric CM.

The grip conditions were counterbalanced among all subjects with half completing session 1 with the *2 Digit* → *5 Digit* grip condition and half completing the session with the *5 Digit* → *2 Digit* grip condition. For those beginning with the *2 Digit* → *5 Digit* grip condition in session 1 the first block of lifts began with completing 10 consecutive “neutral” practice trials with 2 digits to allow the participant to familiarize themselves with the task, texture, and full weight of the object with the CM located in the center. The next block included 10 consecutive trials with the CM on the left side of the device (*2 Digit* L_CM_), immediately followed by 10 transfer trials that were completed with 5 digits on the same CM condition (*5 Digit* L_CM_) after the verbal cue of “switch to 5 digits” was given. Similar to previous studies, each participant completed another block of 10 neutral trials with two digits to wash out any positive or negative learning transfer from the previous block (Fu et al., 2011). The final block included the change in CM location (R_CM_) with 10 consecutive trials with two digits (*2 Digit* R_CM_), followed by the transfer to 10 five-digit trials for the right CM condition (*5 Digit* R_CM_). For session 2 the procedure was repeated, however the order of grip-type conditions was opposite to that experienced in session 1 (i.e., 10 neutral (*5 Digit*) trials, 10 (*5 Digit* L_CM_) trials to 10 (*2 Digit* L_CM_) trials, 10 neutral (*5 Digit*) trials, and 10 (*5 Digit* R_CM_) trials to 10 (*2 Digit* R_CM_) trials).

### Data Processing

Forces and torques were recorded at each object side using the force transducers mounted under the grip surface and the position of the device was recorded by the electromagnetic sensor using custom written software in WinSC/Zoom (Umeå University, Sweden). The outcome measures are defined as:

1. Peak object roll: the angle of the object in the frontal plane, recorded within 250 ms after lift onset. This indicates the subjects’ ability to accomplish the task goal (object roll minimization). Positive values represent counterclockwise roll (towards the left) and negative values represent clockwise roll (towards the right).
2. Load force (LF): tangential component of the force produced by the digits measured in Newtons (N).
  a. LF difference = LF left – LF right. Positive values indicate a larger LF from the thumb on the left grip surface, and negative values indicate larger LF from the finger(s) on the right grip surface.

3. Grip Force (GF): average normal component of the force produced by the digits measured in Newton (N).
4. Center of pressure (CoP): vertical coordinate of the point of application of the digits on the grip surface-measured in (cm).
  a. CoP Difference = CoP left – CoP right. Positive values indicate higher thumb placement than finger(s) CoP, whereas negative values indicate higher finger(s) placement than thumb CoP.
5. Compensatory Torque (Tcom): anticipatory torque generated by the hand, in response to object torque, measured in Newton centimeters (Ncm). This was computed using a formula from a previous study (Lee-Miller et al., 2021).

### Statistical Analysis

A repeated measures ANOVA was used to examine the transfer of learning between two and five digits for each CM location. The Greenhouse-Geisser was applied to adjust the degrees of freedom. Post-hoc statistical comparisons were conducted using a Bonferroni correction with a *p* ≤ 0.016 significance level. Comparisons were made between the novel (trial 1) and pre-transfer (trial 10), the pre-transfer and transfer trial (trial 11) and transfer trial and the final trial (trial 20). These comparisons allowed us to determine whether learning was achieved through consecutive lifts when using an initial grip type for all CM locations. All statistical analyses were performed using SPSS software (SPSS 27.0, Chicago, IL, USA).

## RESULTS

The aim of the study was to investigate whether learned grasping manipulations are transferable across different grasp configurations or constrained to the specific conditions under which they were learned. Fig. 2 displays the GF, LF, COP, Tcom and roll traces for a representative subject during a novel (1^st^) and learned (10^th^) trial using a two-digit grip, and data from the transfer (11^th^) and final (20^th^) trials using a five-digit grip with the CM located on the object’s left side. In the novel trial, the subject has not previously experienced the object with the off-center CM and likely relies on the object’s (symmetrical) visual cues to generate the Tcom (second row, red horizontal line). As a result, the LF and CoP of the thumb and index finger are generally similar at lift onset, and the resulting Tcom falls well below the target Tcom to prevent roll (Fig. 2, vertical dashed line). Thus, the object rolls ∼10 degrees towards the finger side at lift onset before a corrective response is initiated. By the learned (10^th^) trial, the subject modulated their digit placement (CoP) by moving their thumb slightly higher than their finger on the device while increasing their LF difference between the thumb and digits in order to generate a Tcom of the correct magnitude and direction to minimize object roll.

**Fig 2.**
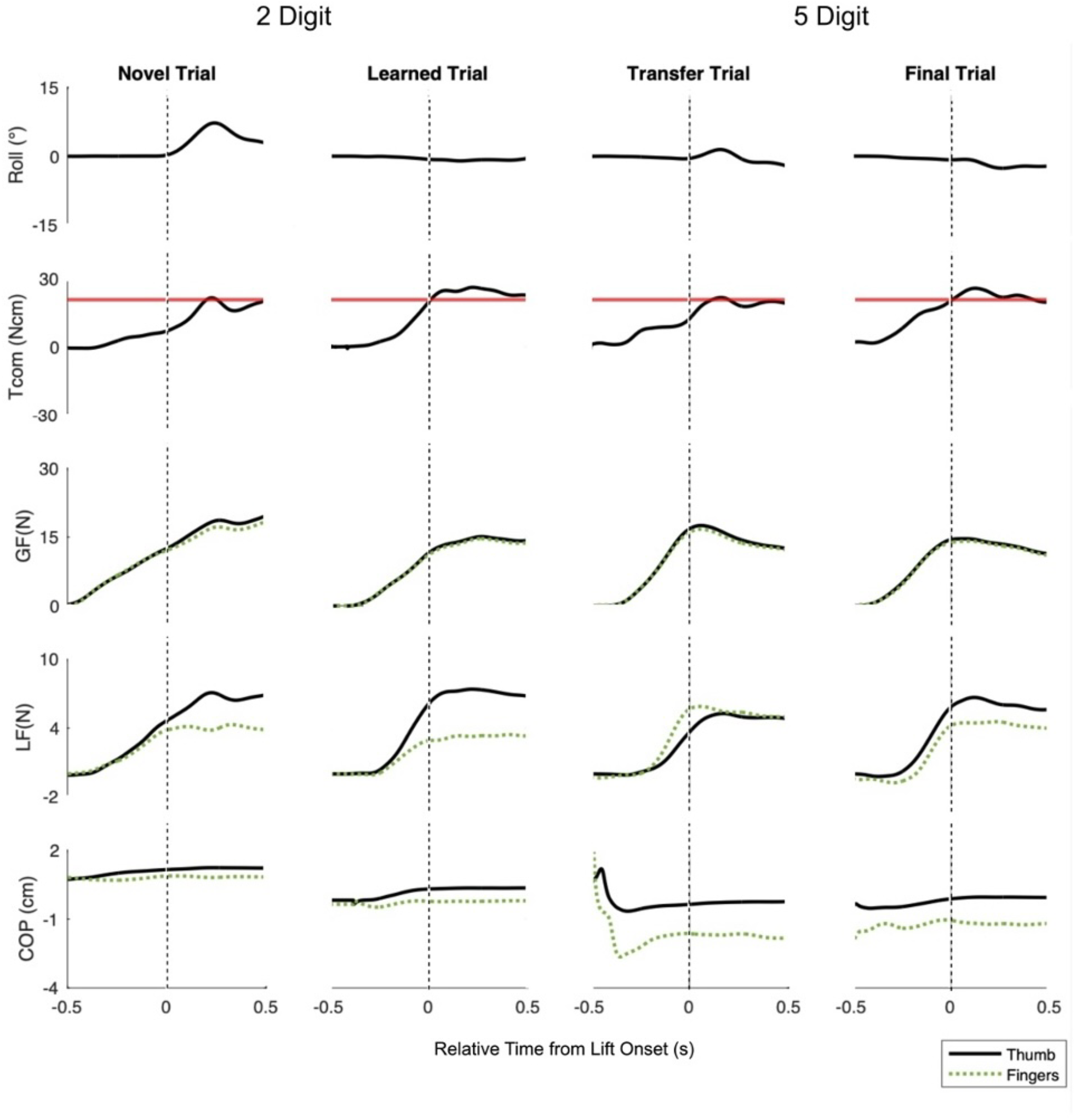
Representative trace plots for the left center of mass (CM). Traces for the novel, learned, transfer, and final trials of all measures for 2-digit and 5-digit grasps. Tcom = compensatory torque, GF = grip force, LF = load force, COP = center of pressure. Vertical dashed lines represent time at lift onset.

On the transfer (11^th^) trial, the subject was asked to switch their grip to using all 5 digits. Although the LF was no-longer modulated at lift onset, the CoP still reflected the object’s CM location leading to a Tcom in the appropriate direction, and thus only a slight roll after lift-onset. Thus, the change in grip type elicited a strategy among CoP where subjects placed their thumb much higher than their digits and evenly distributed their LFs compared to the pre-transfer (10^th^) trial. On the final trial using 5 digits, both the LF and CoP reflect the CM location, with the Tcom reflecting the target and minimal roll.

Similar to previous studies (see Fu et al. 2010; Zhang et al., 2010), by trial 3 performance was stable in comparison to trial 10 (p > .05 for all conditions). Fig. 3a shows the Tcom for the novel trial, pre-transfer trial, transfer trial, and final trial for the left and right CM across all subjects for both the two and five-digit grasp conditions and both CM locations. Across all conditions, there was a significant effect of trial (novel, learned, and transfer) on both (left, right) CM locations. This indicates that the manipulation strategies were influenced by the switch in grip conditions (*2 Digit* → *5 Digit* and *5 Digit* → *2 Digit*), with consistent adjustments observed across trials; 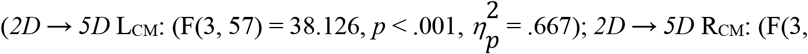 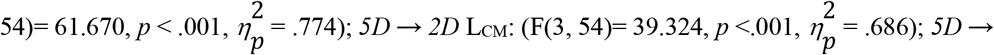 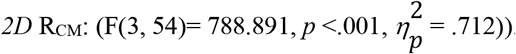. Post-hoc tests showed that the Tcom was significantly higher for the pre-transfer (10^th^) trial than the novel (1^st^) trial for both left and right CM locations and grip conditions (*2D→5D* (L_CM_), *2D→5D* (R_CM_), *5D→2D* (L_CM_), and *5D→2D* (R_CM_)). On the transfer (11^th^) trials, the Tcoms were significantly lower than the pre-transfer (10^th^) only for the *5 Digit* → *2 Digit* condition, left CM. For the novel (1^st^) and transfer (11^th^) trials, post-hoc comparisons showed that for both *2 Digit* → *5 Digit* and *5 Digit* → *2 Digit* conditions, and both (right, left) CM locations, the Tcom for the novel (1^st^) trial was significantly lower than the transfer (11^th^) trials in all cases. For the transfer to final lifts of the same condition, the Tcoms were statistically indistinguishable except for the *2 Digit* → *5 Digit* condition where Tcom was significantly lower in the transfer (11^th^) trial compared to the final (20^th^) trial for the right CM).

**Fig 3.**
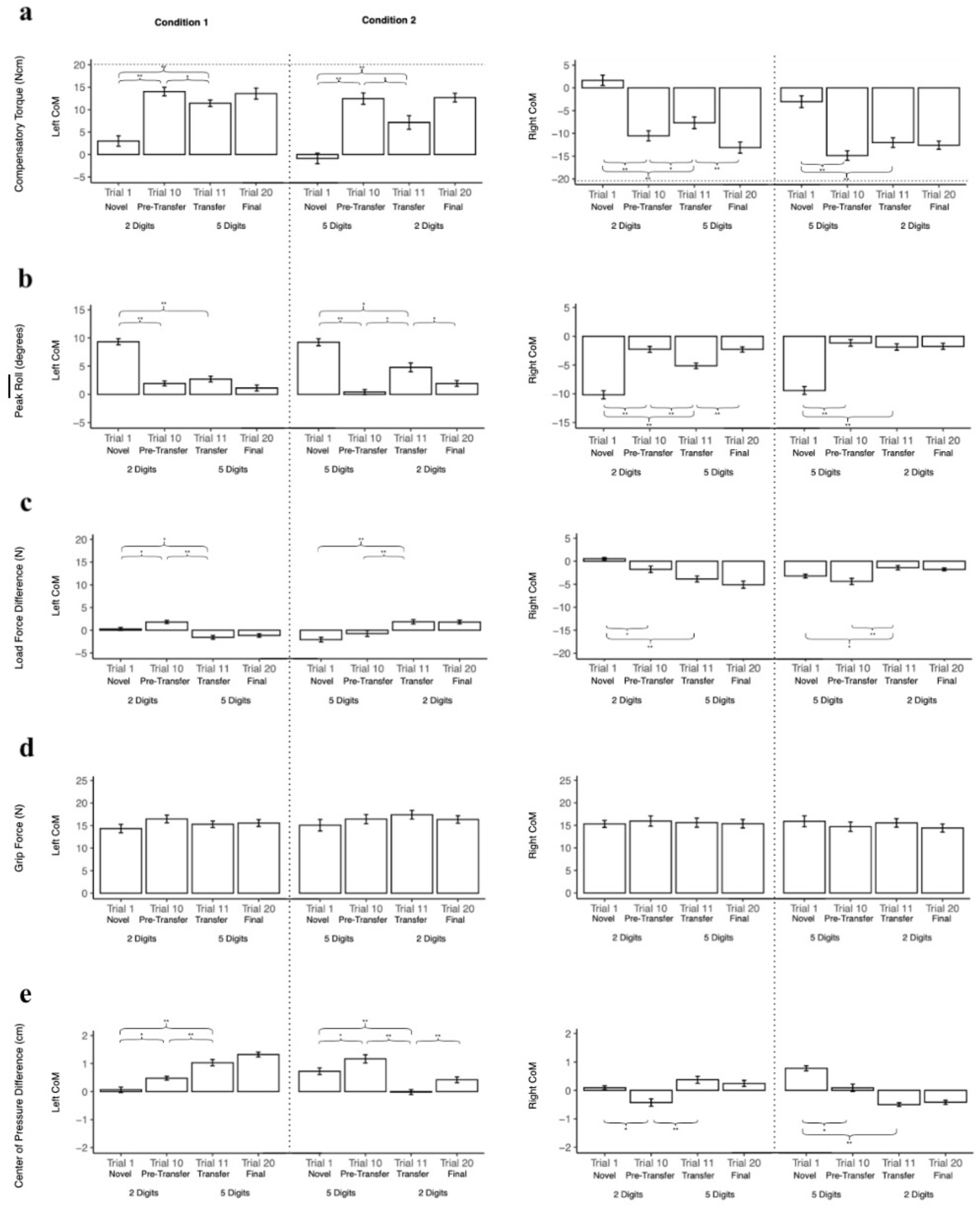
a) Compensatory torque (Tcom) (A) and b) Roll across conditions. Group means (± S.E.M) for the left CM and right CM for the novel trial 1, pre-transfer trial 10, transfer trial 11, and final trial 20. Horizontal dashed lines indicate target Tcom in a. The vertical dotted lines separate *2 Digit* → *5 Digit* Condition (left panel) with *5 Digit* → *2 Digit* Condition (right panel). Asterisks denote significant differences between trials.

The results for the peak object roll generally mirrored the results for the Tcom (Fig. 3b). There was a significant effect for all trials for both (right, left) CM locations, which suggest consistent adjustments in object roll control, reflecting a learned manipulation strategy; 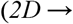 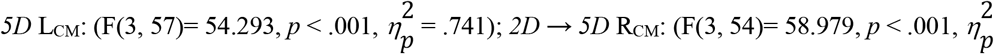 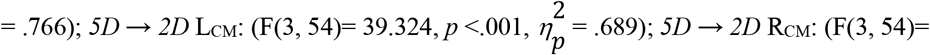 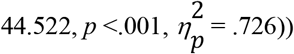.Post-hoc tests shows that there was significantly less object roll in the pre-transfer (10^th^) trial than the novel (1^st^) for both grip conditions and both CM locations (*2 Digit* → *5 Digit* (L_CM_), *2 Digit* → *5 Digit* (R_CM_), *5 Digit →2 Digit* (L_CM_), and *5 Digit* → *2 Digit* (R_CM_)) in all cases (See Fig. 3B). The post-hoc comparisons showed that the transfer (11^th^) trial displayed significantly more roll compared to the pre-transfer (10^th^) trial for *2 Digit* → *5 Digit* condition, right and for *5 Digit* → *2 Digit* condition, left CM. Between the novel (1^st^) and transfer (11^th^) trials there was significantly less roll for *2 Digit* → *5 Digit* condition at both (left, right) CM locations, and *5 Digit* → *2 Digit* condition, right CM and also rolled significantly less at *5 Digit* → *2 Digit* condition, left CM. In the final (20^th^) trial, the object rolled significantly less for *2 Digit* → *5 Digit* condition, right CM) and in the *5 Digit* → *2 Digit* condition, left CM.

Tcom is generated by a combination of load and grip force modulation and digit placement, and peak object roll is inversely related to the extent to which Tcom matches the object’s external torque. Subjects modulated the LF difference between object sides for all trials and CM locations (significant main effect) 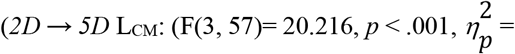 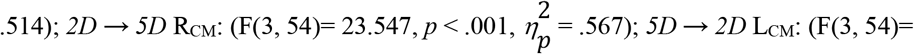 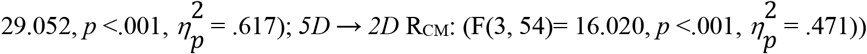. As seen in Fig. 4a, the post-hoc tests showed that LF differences were significantly larger on the thumb side in the pre-transfer (10^th^) trials for the *2 Digit* → *5 Digit* condition for both (right, left) CM locations. The transfer (11^th^) trial the LF was significantly larger on the finger side compared to the pre-transfer (10^th^) trials for the *2 Digit* → *5 Digit* condition, left CM and *5 Digit* → *2 Digit* condition for both (left, right) CM locations. Between the novel (1^st^) and transfer (11^th^) trials, *2 Digit* → *5 Digit* condition was significantly greater on the finger side for the left CM, and also for the right CM. For the *5 Digit* → *2 Digit* condition, the novel (1^st^) trial was significantly larger on the finger side for the left CM and for the right CM compared to the transfer trials.

In contrast to LF, there were no significant main effects of grip type (*p* > .05) on GF across each CM location or trial or between the two object sides (Fig. 4b) for the GFs. This indicates that each subject exerted similar net GFs among all digits used for both grasp conditions.

Subjects modulated digit location to the CM as seen by significantly different vertical separations between digit center of pressure (CoP) for all trials and CM locations 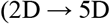 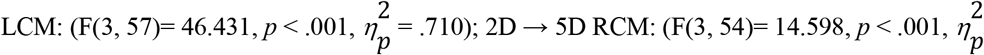 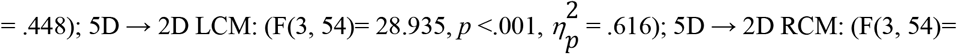 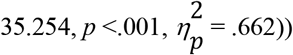. Post-hoc tests showed that the pre-transfer (10^th^) trial had a significantly higher CoP on the thumb side than the novel (1^st^) trial for *2 Digit* → *5 Digit* condition, for both the left CM location, and higher CoP on the finger side for the right CM. For the *5 Digit* → *2 Digit* condition, the novel (1^st^) trial was significantly higher on the thumb side for the right CM, while the pre-transfer (10^th^) trial was significantly higher on the thumb side for the left CM) (see Fig. 4c). The transfer (11^th^) trial was significantly higher on the thumb side than the pre-transfer (10^th^) trial in *2 Digit* → *5 Digit* condition, left CM. The right CM for *2 Digit* → *5 Digit* revealed a higher CoP difference on the thumb side on the transfer (11^th^) trial than the pre-transfer (10^th^) trial. In *5 Digit* → *2 Digit*, only the left CM was significantly higher on the thumb side in the pre-transfer (10^th^) trial compared to the transfer (11^th^) trial. Between the novel and transfer trials, the CoP difference was higher on the thumb side for the transfer (11^th^) trial for *2 Digit* → *5 Digit*, left CM. For *5 Digit* → *2 Digit*, the novel (1^st^) trial was significantly higher on the thumb side compared to the transfer (11^th^) trial for both (right, left) CM locations. For the transfer and final trials, the CoP difference was also significantly higher on the thumb side for the final (20^th^) trial compared to the transfer (11^th^) trial for *5 Digit* → *2 Digit*, left CM.

## DISCUSSION

The results of this study highlight the effects of different grip configurations (2 Digit vs. 5 Digit) and center of mass (CM) locations on the extent to which learned object manipulation can be transferred. For our task, subjects were required to lift an asymmetrical object while minimizing roll caused by an external torque. The study explored transitions between these grip types across novel, learned, transfer, and final trials. The results indicated that subjects were able to generate a torque in the correct direction in the new digit configuration but required additional experience using that configuration to refine its specific digit positions and forces and minimize roll further. This suggests that object manipulation is learned, at least partially, through high-level (task) representation, rather being exclusively dependent on the effectors used to learn the manipulation.

In order to determine whether transfer was achieved, comparisons were made between the novel (1^st^) trial and the transfer (11^th^) trial and the pre-transfer (10^th^) trial and the transfer trial. For evidence of transfer, Tcom needs to be maintained between the pre-transfer (10^th^) and the transfer (11^th^) trial. Transfer would also be indicated by a significant improvement between the novel (1^st^) trial and the transfer (11^th^) trial, showing that subjects maintained some features of the learned behavior within the post-transfer grasp that are functionally better than the novel trial (Fu et al., 2011). The following are possible scenarios subjects displayed: (a) no transfer (trial 11 is significantly different from trial 10 but not trial 1); (b) imperfect transfer (trial 11 is significantly different than trial 1, but still different than trial 10); and (c) perfect transfer (trial 11 is not significantly different than trial 10). Based on previous findings (Fu et al., 2011), a lack of difference between trials 10 and 11 would the strongest evidence of transfer. The results suggest that subjects initially struggled to keep the object steady with the novel grip configurations (i.e., novel [1^st^] trial) as evidenced by the insufficient Tcom and highest object roll in the novel trial. However, as they progressed to the pre-transfer (10^th^) trial, significant increases in Tcom and reductions in object roll indicated subjects’ learning and adaption. These findings were consistent with previous studies of implicit learning that subjects are able to minimize object roll within blocked conditions (Fu et al., 2010; J. R. Lukos et al., 2008). The most notable finding in our study was that subjects’ performance in the transfer (11^th^) trial was significantly better than the novel (1^st^) trial in the correct direction, for all grip conditions and CM locations. This indicated that subjects were able to transfer some of the information learned from one grip type to the other without disruption to the overall goal of the task (Budgeon et al., 2008; Fu et al., 2011; Santello and Soechting, 2000). However, during the transfer (11^th^) trials when the immediate switch in grip type configuration occurs, there was a reduction in Tcom in 1 out of the 4 conditions and an increase in object roll in half the conditions compared to pre-transfer trials, indicating there was not a “perfect transfer” between 2-digits and 5-digits in many cases. This demonstrates the change from one grip type to another introduced challenges in object manipulation, which initially destabilized object control. Unlike the Fu et al. 2011 study that investigated simpler grip configuration transfers (*2-to 3-Digit and 3-to 2-Digit*), the whole-hand grasp condition we used resulted in a larger lever arm than simply 2 or 3 digit grasps, and thus a force sharing pattern among 4 fingers to counter the thumb), with the change in degrees of freedom further altering the complexity.

The LF differences between the thumb and finger sides of the object were evident, demonstrating that subjects distributed their GF to stabilize the object. During the *2 Digit* → *5 Digit* condition, in the pre-transfer (10^th^) trial when the CM location was on the left (i.e., more weight loaded on the thumb), the LF was higher on the thumb side. In precision grasping, the thumb plays a critical role in stabilizing objects, especially in tasks where precision and load balancing are necessary (Westling and Johansson, 1984, p. 1988; Zatsiorsky et al., 2004). However, in the transfer (11^th^) trial, the LF shifts more towards the fingers, indicating that subjects had adjusted their force distribution as they transferred to the different grip configuration. This is remains evident when object CM remains on the left, but the grip switch pre-transfer (10^th^) from *5 Digit* → *2 Digit* LF remains on the finger side before transferring to the thumb side in the transfer (11^th^) trial, indicating that subjects were modulating their digit forces in accordance with studies of multi-digit interactions and force-sharing patterns in whole-hand grasping. The addition of digits opposing the thumb involve utilization of a force sharing patterns, with multiple combinations of force opposing the thumb being equally viable solutions to prevent tilt (e.g., Albert et al., 2009; McIsaac et al., 2009; Rearick and Santello, 2002; Zhang et al., 2011). For the right CM condition, regardless of the transfer grip-type condition (i.e., *2 Digit* → *5 Digit* or *5 Digit* → *2 Digit*), results indicated that participants distributed more LF on the finger side under the heavier side of the object, using the thumb as a stabilizer. This allowed for transfer of Tcom and minimized object tilting (Fu et al., 2010).

With regard to center of pressure (CoP), our results revealed important adaptations in digit positioning relative to the object’s CM. We observed significant variations in the CoP difference across all trials, indicating that subjects created different vertical separations between the thumb and fingers after switching between precision and whole-hand grasping, similar to previous findings among transfer between different grip types (see Fu et al., 2011; *2-digit and 3-digit* grip configurations).

When lifting with two digits, subjects demonstrated less of a difference between the thumb and finger CoP, working together as a unit to stabilize the object. Both digits adjusted their force output in a coordinated manner to counteract the object’s uneven weight distribution. Results also revealed that the thumb and fingers shared the responsibility for generating stabilizing forces, while subjects managed to control the four digits as one unit to generate Tcom.

This positioning helps distribute forces effectively, maintaining a more stable CoP across the hand.

It is important to note that our analysis focused on the CoP difference between the thumb and finger sides respectively. While this approach highlights the overall change in the digit positioning, future work should explore the thumb and finger CoP’s independently to provide additional insights into the specific contributions of each digit to grip stability and object manipulation.

Subjects had the ability to utilize various solutions to achieve the goal of our task. Our results show that subjects successfully modulated their LFs in order to minimize roll when lifting an asymmetrical object. We only saw a partial solution of higher digit placement in the left CM condition regardless of the addition or removal of effectors. Thus, the imperfect transfer may have been due to the differential transfer of digit positions and force, a finding that is consistent with those reported by Zhang et al. (2010) (see Introduction). A limitation of this study was the inability to measure the force of each individual finger separately. This constraint prevented us from determining the specific contributions of individual digits to the overall observed force and coordination patterns. Without finger-specific data, it presents a challenge to assess how forces are redistributed among the four fingers during task execution. Such redistribution is critical for understanding the strategies employed to achieve task goals, particularly in complex manipulative tasks where fine-tuned force modulation is essential. Although we were able to analyze the overall effect, it would be beneficial for future studies to assess the redistribution of forces among the four fingers reducing the need of modulating digit positions (Marneweck and Grafton, 2020). Another limitation is that while we counterbalanced between the transfer directions, we did not between the CM locations. As in previous studies, however, there did not appear to be a systematic difference in the ability to transfer between CM locations across variables.

An open question is how effector (digit)-based sensory inputs from touch and proprioception are used to enable a high-level representation (Tcom) to be exerted when using a grip configuration that differs from the configuration used to learn the task. Although our study cannot answer this question, the use of a high-level representation regardless of grip configuration would require mechanisms that, ultimately, allow the comparison of Tcom using a new grip configuration relative to the Tcom learned with a different grip configuration. Such comparison, in turn, requires sensory inputs acquired through different grip configuration to be integrated in a way that the CNS can interpret as their combined effect (i.e., target Tcom). Future work should address the sensorimotor transformations involved with error-free and partial (current results) transfer of learned high-level task representations.

Unlike previous studies, our study demonstrates how challenging grip transitions, asymmetrical object dynamics, and CM location interact to reveal both the strengths and limitations of learned manipulation transfer. By showing that subjects can imperfectly transfer learned Tcom strategies whole also requiring local adjustments in load force and digit positions, our results suggest an important interaction between high-level (task) representations, and effector-specific adaptations. This complexity extends beyond simpler 2-to 3-digit studies, offering unique insights into the mechanisms underlying dexterous object manipulation.

The ability to grasp objects with two digits and whole-hand is crucial for human dexterity and for the performance of a wide range of everyday tasks. This raises important questions about whether people with various neurological conditions are able to generalize behaviors from one learned condition to another and demonstrate the use of higher-level (task) representations. This is particularly relevant for understanding how impairments in motor planning and execution may affect the ability to transfer learned skills across different contexts, potentially offering insights into targeted rehabilitation strategies. The present study demonstrated that learned object manipulations can be transferred and refined with practice, regardless of adding or removing digits. Our findings further suggest the existence of high-level, effector-independent representation object manipulations that can be learned and transferred.

## Acknowledgements

This paper was supported by grants to AMG and MS by the National Science Foundation grants BCS-1827725 and BCS-1827752.

## Figure Legends

**Fig 4. Components of Tcom across conditions (a-d).** Group means (± S.E.M.) for **(a)** load force difference, **(b)** Grip Force, and **(c)** center of pressure difference for the left CM (top panel) and right CM (bottom panel) for the novel trial 1, pre-transfer trial 10, transfer trial 11, and final trial 20. The vertical dotted lines separate *2 Digit* → *5 Digit* Condition (left panel) with *5 Digit* → *2 Digit* Condition (right panel). LF = load force, GF = grip force,, COP = center of pressure. Asterisks denote significant differences between trials. Asterisks denote significant differences between trials.

